# Extrachromosomal circular DNA, microDNA, without canonical promoters produce short regulatory RNAs that suppress gene expression

**DOI:** 10.1101/535831

**Authors:** Teressa Paulsen, Yoshiyuki Shibata, Pankaj Kumar, Laura Dillon, Anindya Dutta

**Affiliations:** Department of Biochemistry and Molecular Genetics, University of Virginia School of Medicine, Charlottesville, VA, 22908, USA

## Abstract

Interest in extrachromosomal circular DNA (eccDNA) molecules has increased recently because of their widespread presence in normal cells across every species ranging from yeast to humans, their increased levels in cancer cells, and their overlap with oncogenic and drug-resistant genes. However, the majority of eccDNA (microDNA) are too small to carry protein coding genes. We have tested functional capabilities of microDNA, by creating artificial microDNA molecules mimicking known microDNA sequences and have discovered that they express functional small regulatory RNA including microRNA and novel si-like RNA. MicroDNA is transcribed *in vitro* and *in vivo* independent of a canonical promoter sequence. MicroDNA which carry miRNA genes form transcripts which are processed into mature miRNA molecules, through the endogenous RNA-interference pathway, which repress a luciferase reporter gene as well as endogenous mRNA targets of the miRNA. Further, microDNA containing sequences of exons repress the endogenous gene from which the microDNA was derived through the formation of novel si-like RNA. We also show that endogenous microDNA associate with RNA polymerases subunits POLR2H and POLR3F. Together, these results suggest that microDNA may modulate gene expression through the production of both known and novel regulatory small RNA.

## INTRODUCTION

Extrachromosomal circular DNA (eccDNA) exists within all eukaryotic organisms tested (1,2,11–17,3–10) and come from consistent hotspots within the genome including 5’UTRs, exons, and CpG islands (13–17). For a recent review see (18). The majority of eccDNA in normal cells are small in size, 200-400 base pairs, though they can range up to tens of thousands of base pairs (13,14,16,17,19). Mega-base sized eccDNA molecules have been known for some time as double minutes that amplify oncogenes, and smaller forms of eccDNA (in the tens of kilobase range) have recently been found to also amplify oncogenes (20–22), drug-resistant genes (23), and tissue specific genes (13,16,19,24). The smallest type of naturally occurring eccDNA, <500 bp, are called microDNA (13,14,16). The limited size of these molecules precludes them from carrying full protein coding gene sequences and or promoter sequences.

By electron microscopy, endogenous microDNA are both single and double-stranded (13,16). We hypothesized that microDNA can be transcribed based on previous research that suggested that single stranded circular DNA of 34-89 base pairs can be transcribed without a promoter *in vitro* and within cells by a rolling circle mechanism in which the polymerase travels around the circle 12-260 times (25–27). However these circles are much smaller than the average microDNA. *Paramecium tetraurelia*, on the other hand, creates double-stranded DNA circles of unknown length by fusing transposon derived sequences and these circles also express small regulatory RNA (28).

Here we investigate whether microDNA-mimics are capable of being transcribed in mammalian cells without a canonical promoter and whether the transcripts are functional within a cell. Both single stranded and double stranded microDNA (ranging from ~180-400 base pairs) are transcribed without a promoter *in vitro* and *in vivo*. RNA is produced from both strands of microDNA without strand bias. MicroDNA containing miRNA coding sequences, but without the promoter of the gene, produce functional miRNA capable of knocking down both a luciferase reporter and endogenous mRNA targets. MicroDNA is known to be enriched in genic regions (16) so that some microDNA carry exon sequences. We report that microDNA arising from exons can also affect gene expression by expressing novel si-RNA that targets the parental gene that the microDNA was derived from. We also show RNA polymerase subunits are associated with endogenous microDNA, giving further evidence that microDNA molecules can be transcribed in cells. Together these results show that microDNA could produce functional regulatory RNA, both miRNA and novel si-RNA, and suggest a new mechanism of how genomic plasticity and instability can lead to changes in gene expression.

## MATERIALS AND METHODS

### Cell Culture

HCT116, 293A and 293T cells were cultured in Dulbecco’s modified Eagle’s medium (DMEM) supplemented with 10% fetal bovine serum, 100 U/ml penicillin and 100 μg/ml streptomycin in an environment containing 5% CO2 at 37°C. 293T cells were cultured in McCoy’s medium supplemented with 10% fetal bovine serum, 100 U/ml penicillin and 100 μg/ml streptomycin in an environment containing 5% CO2 at 37°C.

### Artificial microDNA synthesis

Artificial microDNA molecules containing known microDNA sequences were created using a protocol published by Du Q, et al (29). In short, microDNA sequences were amplified out of HeLa genomic DNA using PCR. A circularly permuted molecule was created through PCR amplification of each half, which were then cloned in the appropriate order into a pUC19 plasmid using an In Fusion HD Cloning Kit (Takara). The substrates were amplified using Phusion Polymerase PCR (NEB). The linear double-stranded molecules were denatured and renatured to produce both the parental linear molecules and a circle with nicks on each strand nearly half-way around the circle which were then ligated by Taq Ligase (NEB). The products were taken through 10 cycles of denaturation, annealing and ligation to enrich for circular DNA. Residual linear DNA was digested using ExoI and ExoIII (NEB) and the products separated on a denaturing PAGE gel. The band corresponding to dsDNA circles was excised and the DNA was extracted for the *in vitro* transcription reactions. Unless otherwise noted, the final products after the ligation cycles and exonuclease digestion was transfected into cells.

### *In vitro* transcription assay

The artificial microDNA (100-200 ng per 50 μL reaction) was transcribed for 4 hours *in vitro* in an IVT buffer (40 mM Tris-HCl pH 7.9, 6 mM MgCl2, 10 mM DTT, 2 mM spermidine, 0.1 mM NaCl), rNTPs (2 mM rATP, 2 mM rCTP, 2 mM rGTP, 0.4 mM rUTP), [α-32P]-UTP (volume dependent on radioactivity), and 20 μg of HeLa nuclear extract (Millipore) at 37°C. The radioactive product RNA was heat denatured and run on a denaturing (urea) PAGE gel. The gel image was captured using a phosphor imaging screen and a gel imager.

### Transfections of microDNA

MicroDNA was transfected in cells using Lipofectamine LTX according to the manufacturer’s instructions. RNA was isolated 24 hours after transfection. Plates with 0 ng of microDNA received 100 ng of a GFP plasmid as carrier. By transfection of a GFP expressing plasmid we estimate that 70-80% of the 293 cells take up the plasmid, but we are unable to say what fraction of the input plasmid enters the cell or the nucleus.

### RNA Isolation and Quantification

RNA was extracted using TRIZOL according to the manufacturer’s instructions (Ambion). The cDNA was created using the miScript II RT kit (QIAGEN). Specifically, the pre-microRNA sequences were quantified by creating cDNA with the miScript II RT Kit with miScript HiFlex Buffer and then amplified by QPCR with primers that flank the mature microRNA sequence within the pre-microRNA molecule. The mature microRNA sequences were quantified by creating cDNA of mature microRNA with the miScript II RT Kit with miScript HiSpec Buffer and then amplified by QPCR using a primer which targets the microRNA sequence and the 10X miScript Universal Primer. The miScript II kit is to selectively amplify the short microRNA and not the longer product that may be created by ligation of the 3’ adaptor to pre-microRNA. QPCR was performed using Power SYBER Green Master Mix (Life Technologies).

### Luciferase assays

DNA oligonucleotides designed to carry the sequence of the mature miRNAs encoded by the microDNA (miR191, miR126, miR145) were cloned into the siCHECK (Promega) vector into the 3’UTR of the Renilla luciferase gene such that the sequence complementary to the miRNA was in the + strand. The effect of miRNA produced by microDNA on the siCHECK vector was quantified using the Dual-Luciferase Reporter Assay System (Promega) according to manufacturer’s instructions.

To test endogenous microDNA repression on a luciferase reporter 20 ng of either the control siCHECK vector and the siCHECK vector containing the sequence homologous to a portion of the TESC-AS1 microDNA (listed in the supplemental figures) were transfected into a 24-well plate. After 24 hours the luminescence of the samples was quantified using the Dual-Luciferase Reporter Assay System (Promega) according to manufacturer’s instructions. The specific sequence cloned into the siCHECK vector homologous to the TESC-AS1 microDNA: GCTCTCCTGAAGGCTCTGCAG.

### Construction and infection of HaloTag vectors

The plasmid pENTR4-HaloTag which encoded the HaloTag sequence was obtained from Addgene. POLR2H and POLR3F cDNA were amplified from hORFeome V5.1 clones. PCR amplified POLR2H or POLR3F cDNA were inserted in frame downstream of HaloTag sequence. HaloTag, HaloTag-POLR2H or HaloTag-POLR3F were subcloned into the pCW plasmid vector.

293T cells were transfected with plasmids pCW-HaloTag (or pCW-Halo-fusions), psPAX2 and pCMV-VSV-G using lipofectamine 2000. Lentivirus was harvested from the supernatant after 48 hours, cleared by centrifugation and passed through a 0.45 μm filter. To obtain stably transduced 293A clonal cell derivatives expressing HaloTag fusion proteins, lentivirus was added to 293A cells in the presence of 6 μg/ml polybrene followed by selection with 3 μg/ml puromycin, and isolation of clones by dilution cloning.

### HT Cell lysate preparation and pull-down on Halo-Link beads

All the reagents used were pre-chilled and the entire procedure was performed on ice. HT-fusion proteins were induced by adding 1 μg/ml doxycycline to cells in a 15-cm plate. Two days later, cells were washed with PBS, scraped and transferred into a micro-centrifuge tube. After centrifugation five packed cell volume (PCV) Hypotonic Buffer was added to cell pellets, allowing the cells to swell, for 15 min on ice. Hypotonic Buffer was removed after spinning. Cells were suspended in 0.5 PCV of Buffer LS and equal volume of Buffer LS with 600 mM NaCl and 0.2% Triton X-100 was added. Cells were homogenized by passing 10 times through a 27G hypodermic needle and rotated for 20 min. After centrifugation, the supernatant was transfer into a new microcentrifuge tube and equal volume of Buffer LS to the supernatant was added.

Cell lysate was added to the equilibrated HaloLink Resin and incubated by mixing on a tube rotator for 30 min at room temperature. After centrifugation supernatant was saved as sample flow through. Resin was washed with PD Washing Buffer five times. The beads were boiled in Laemmli sample buffer to obtain the eluates.

### MicroDNA extraction and identification

HT or HT-POLR3F/POLR2H associated DNA was purified with QIAprep Spin Miniprep Kit according to its instruction manual and amplified by rolling circle amplification (RCA) as described (14,16).

Initially paired-end high-throughput sequencing (250 cycles PE) was performed on the RCA products according to the manufacturer’s protocol (Illumina) on the Illumina MiSeq at the University of Virginia DNA Sciences Core (Charlottesville, VA, USA) (Table 1). Read quality was checked by program fastqc (FASTX-Toolkit) and was found that the median read quality was less than 28 after position 150 on the read. Therefore to remove the bad quality bases we made each library 150 PE and did the downstream analysis. We used Burrows–Wheeler Aligner with maximal exact matches (BWA-MEM) to align the reads to the human hg38 genome, allowing for split reads under default conditions. Like our previous publication (13,14,30) we used split reads mapped position to identify microDNA co-ordinate at base pair resolution. In summary to identify microDNA we consider paired-end reads that had one end mapping uniquely to the reference genome (mapped end) and the other end not mapping to the reference genome continuously, but coming from a split-read (the junctional read). Furthermore, the two parts of the split read have to flank the linked mapped read. The identified microDNA would represent genomic coordinates of potential microDNA junctions created by the ligation of two ends of a linear DNA. In addition to this we also check polarity (strand information) of both the split read (should be map in the same orientation) and mapped read in pair (this should be opposite to split read).

**Table 1.**
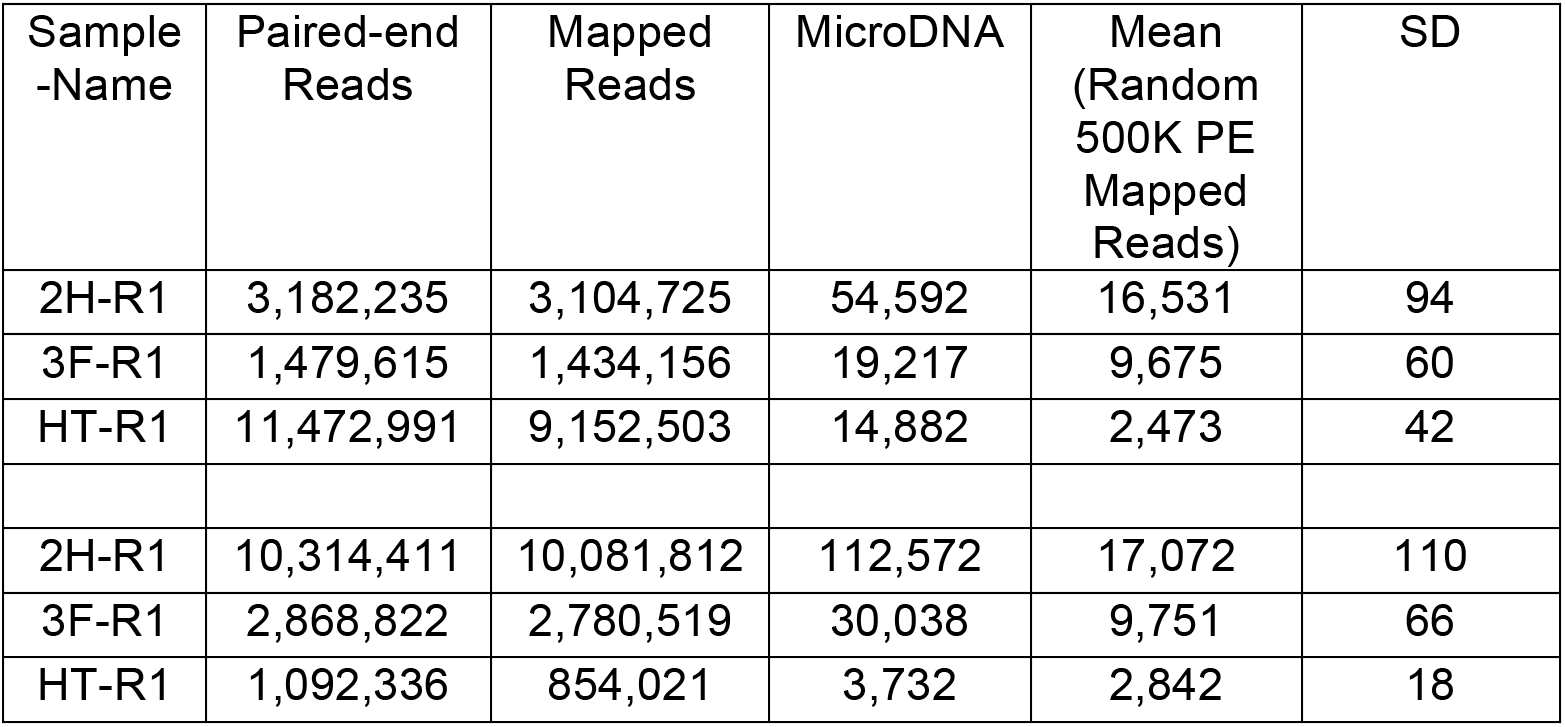
MicroDNA diversity associated with HT, and HT fused with indicated RNA polymerase subunits. Summary of reads obtained from HT, HT-POLR2H and POLR3F associated microDNA libraries (2 independent sequencing runs done on separate days, R1 and R2). Low quality bases from the 3’ end of reads were removed by making 150 (read length) PE reads from 250 (length) PE reads. We also made 75 bases PE reads from the same library to identify microDNA shorter than 150 bp. Total number of unique microDNA (unique microDNA junctions) identified from each run is indicated. To normalize for the number of paired end reads in the three libraries, 500,000 mapped reads were randomly selected from each library and the number of unique microDNAs in the samples determined. This was done 10 times and the mean microDNA number and their standard deviation indicated in the last two columns.

### Evaluation of complexity of microDNA associated with RNA polymerases

The microDNA were identified from the HaLo-Tag pull-downs as above. Because of the differences in number of mapped reads obtained from the HT, HT-POL2H and HT-POL3F associated microDNA, we used a random sampling approach to compare the microDNA complexity among samples (Table 1). The same number of mapped reads was randomly extracted from the HT-POLR2H and HT-POLR3F libraries, as the total number of mapped reads in the HT library. This was done 10 times. Each randomly selected set of reads was processed as above and the number of unique microDNAs identified counted (Table 1). The mean and standard-deviation of the 10 samples is presented in Figure 6C.

## RESULTS

### Synthesis of microDNA mimics

We created synthetic microDNA mimicking known microDNA sequences by utilizing a technique, called ligase assisted mini-circle accumulation (LAMA), which relies on cycles of annealing, ligation and denaturation to produce small DNA circular molecules (Figure 1A) (29). Utilizing sequencing data of microDNA isolated from human cancer cell lines, we designed circles that mimic known microDNA overlapping with microRNA sequences or with exons of non-coding or protein coding genes. Both single stranded and double stranded artificial microDNA molecules were created and isolated (Figure 1B). The sequences of the circular molecules created are listed in Supplemental Table 1. Double-strandedness of specific isoforms was verified by digestion with restriction endonuclease, while topoisomerase I was used to distinguish supercoiled from relaxed circles (Supplemental Figure 1).

**FIGURE 1.**
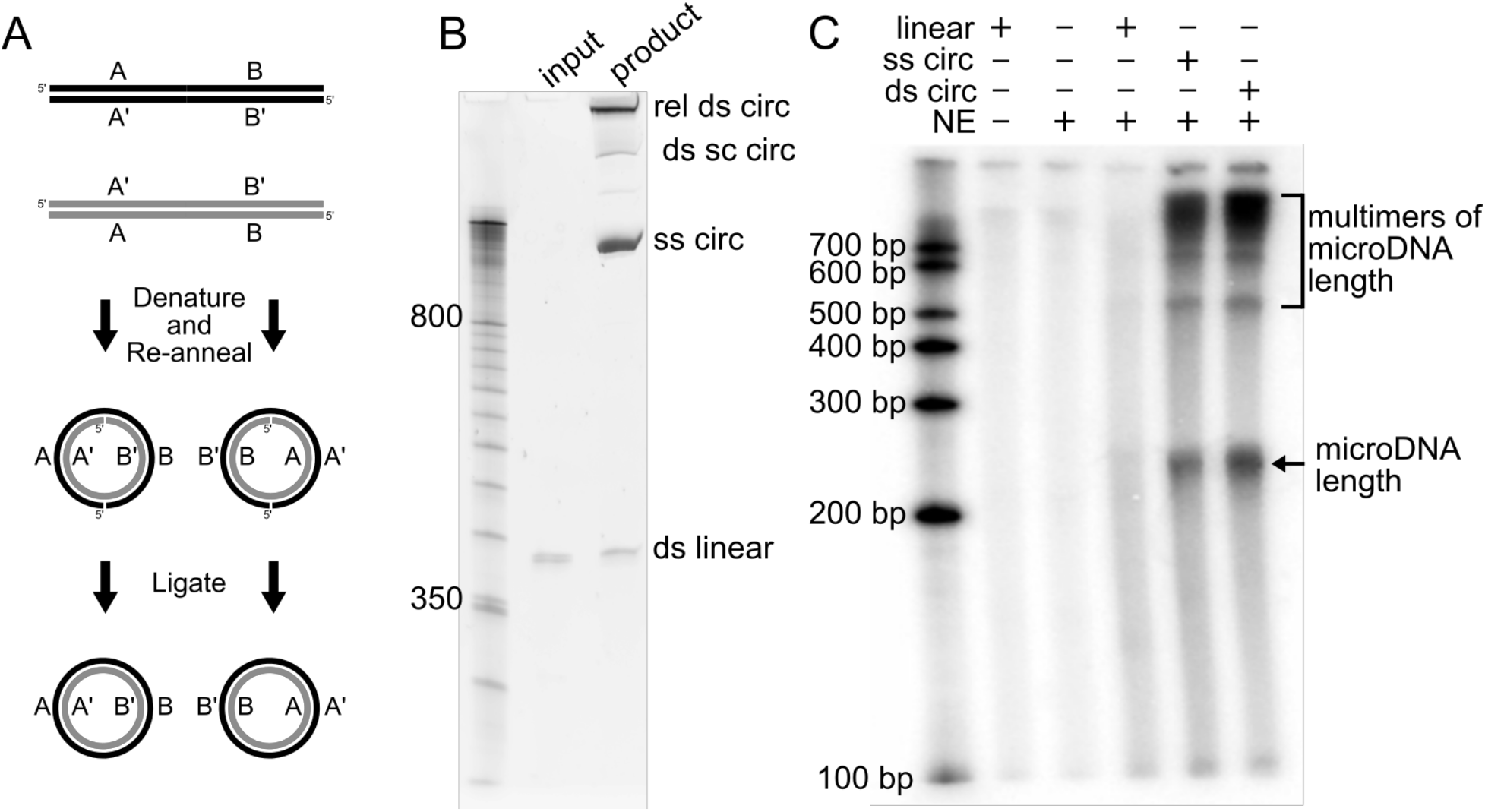
(A) Diagram of artificial microDNA creation by LAMA. (B) Circular and linear products of LAMA run on a denaturing PAGE gel before and after ligation cycles. Circular DNA accumulates to form ssDNA, nicked and supercoiled dsDNA. (C) 32P-UTP labeled RNA run on PAGE gel after in vitro transcription assay. Products are seen at size ranges of multiples of the microDNA length. Ss: single-stranded; ds: double-stranded; sc: supercoiled; rel: relaxed or nicked; circ: circular; NE: HeLa nuclear extract. Representative replicate of duplicates.

### MicroDNA mimics are transcribed *in vitro*

To test whether microDNA are transcribed we used an *in vitro* transcription system using HeLa nuclear extract which contains human RNA polymerases. We isolated the single stranded and double stranded microDNA (containing the hsa-mir-145 sequence) after separation on the denaturing PAGE and added transcriptionally competent HeLa nuclear extract, NTPs, and radiolabeled UTP. Both single stranded and double stranded circular microDNA molecules are transcribed but the linear DNA control containing the same sequence is not transcribed (Figure 1C). Relaxed circles produced significantly more transcripts than supercoiled circles (Supplemental Figure 1). This experiment was repeated with two other microDNA sequences (microDNA carrying hsa-mir-126 sequence and the hsa-let-7a sequence) and similar results were found (Supplemental Figure 2). Each in vitro transcription experiment was validated with a second replicate. The RNA products show distinct lengths that correspond to multiples of the microDNA sequence length. This suggests that some RNA polymerases fall off the DNA template after going around the circle once and some continue around the circle multiple times.

Because the microDNA sequences did not contain known promoters this result also shows that the short double-stranded DNA circles, as well as the single-stranded DNA circles, are transcribed independent of a canonical promoter sequence by human RNA polymerases, whereas a linear DNA fragment is not transcribed. We hypothesize that the bending of the double-stranded DNA enables the binding of TATA-binding proteins (31–34) to recruit RNA polymerase which initiates transcription independent of a canonical promoter.

To determine where transcription initiates on a microDNA mimic (microDNA with hsa-mir-191 sequence), we performed an *in vitro* transcription reaction and then quantified the RNA arising from different regions of the microDNA sequence. We found that the transcription is not uniformly distributed around the circle, but has some sequence bias (Supplemental Figure 3). This suggests that certain sequences within the microDNA are more likely to be bound by transcription initiation machinery than others.

### MicroDNA mimics are transcribed *in vivo*

We next wanted to test whether the artificial microDNA molecules can be transcribed *in vivo*. Because some of the microDNA molecules which have been sequenced from human cancer cells contain miRNA sequences (16), we hypothesized that microDNA may be capable of expressing miRNA (Figure 2A). Artificial microDNA molecules were made that carry miRNA sequences observed in naturally occurring microDNA sequences: hsa-mir-145, hsa-mir-191, hsa-mir-126. The sequences contained only the pre-miRNA part of the gene but not the rest of the primary miRNA, and hence excluded the promoter that is found at the 5’ end of the long primary miRNA, often tens of kb away from the mature miRNA. As the amounts of artificial microDNA are increased, the levels of each of the pre-miRNA transcripts increased by 6-to 30-fold (Figure 2B). At least some of pre-miRNA transcripts are processed into mature miRNAs which are also increased concurrently by 3-to 6-fold (Figure 2C).

**FIGURE 2.**
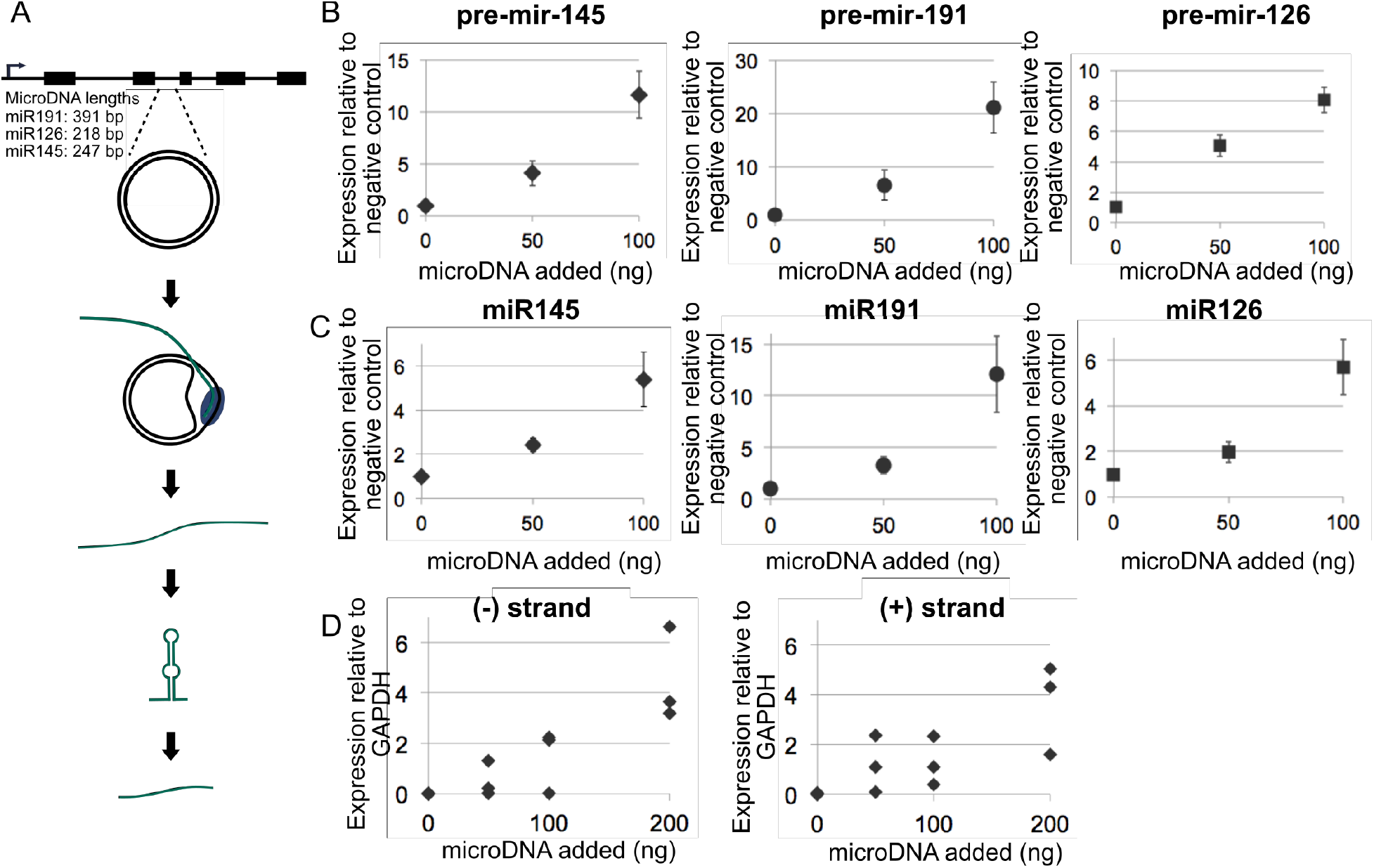
(A) Diagram of transcription of microDNA carrying only the promoterless pre-microRNA part of the microRNA gene. The transcripts are processed by endogenous RNA interference proteins into functional mature miRNA (B) Expression of pre-microRNA molecules after addition of indicated amounts of the corresponding artificial microDNA molecules. Expression is relative to beta-actin (hsa-mir-191 and hsa-mir-126) and GAPDH (hsa-mir-145) and normalized again to the negative control of a transfected GFP plasmid. Mean and S.E. of 3 transfections. (C) Expression of processed mature miRNA molecules after transfection of indicated amounts of artificial microDNA molecules. Expression is relative to beta-actin (hsa-mir-191 and hsa-mir-126) and GAPDH (hsa-mir-145) normalized to negative control. (D) Expression of the + and the – strand of the pre-microRNA, hsa-mir-145, from microDNA molecules relative to GAPDH. Strand-specific primers were used for the reverse-transcription to form cDNA specifically from the (+) or (-) strand of RNA. Results from 3 transfections.

Additionally, the transfection of linear DNA carrying the same sequence as the microDNA does not increase the RNA levels confirming that the DNA must be circularized to be transcribed (Supplemental Figure 4A). To further show that the RNA is arising from circular DNA and not linear contaminants or the genomic sequence, we quantified the RNA using primers that amplify the junction sequence. We see an increase in the RNA spanning the junction as more artificial microDNA is added (Supplemental Figure 4B). The fold increase of the junctional RNA appears significantly higher relative to fold-induction seen with primers known to target the pre-microRNA within the microDNA. We believe this is because there is no endogenous RNA in untransfected cells that spans the junction of the circle. Therefore, the fold-induction using the primers spanning the junction sequence gives a more accurate representation of the quantity of RNA arising from microDNA because there is no endogenous junction-spanning RNA elevating the basal level in untransfected cells.

Because the microDNA is transcribed without a canonical promoter sequence we wondered whether the RNA is equally transcribed from both strands of the microDNA when doublestranded circular microDNA is transfected. Reverse-transcription with strand-specific primers and Q-PCR demonstrated that the microDNA carrying hsa-mir-145 were transcribed relatively equally from either strand (Figure 2D). The relative stabilities of the RNAs arising from the two strands and whether the RNAs are processed by RNA capping and poly-A addition is unknown and will require further research.

Overall, these results suggest that microDNA like molecules can be actively transcribed within cells. The specific character of microDNA allowing for its relative independence from a canonical promoter requires further determination. This research gives insight into how these circular DNA molecules, till now assumed to be inert byproducts of DNA metabolism, could contribute to cell physiology by actively forming RNA transcripts.

### MicroRNA produced from microDNA mimics are functional

To determine whether the microDNA produced functional miRNAs, luciferase reporters containing target sequences complementary to the miRNAs in their 3’ UTRs were co-transfected with the microDNA for dual luciferase assays in 293T cells. Each of the microDNA molecules (containing hsa-mir-145, hsa-mir-191, or hsa-mir-126 sequences) repressed the luciferase reporter carrying the target sequence of the miRNA by >50% after transfection (Figure 3A). Further, the microDNA containing hsa-mir-145 and hsa-mir-191, which carry both the 3p and 5p sequences of the microRNA, are able to repress a luciferase reporter which contains either the 3p or 5p sequence. This shows that the transcripts arising from the microDNA form both mature sequences and function in the same manner as endogenous microRNA.

**FIGURE 3.**
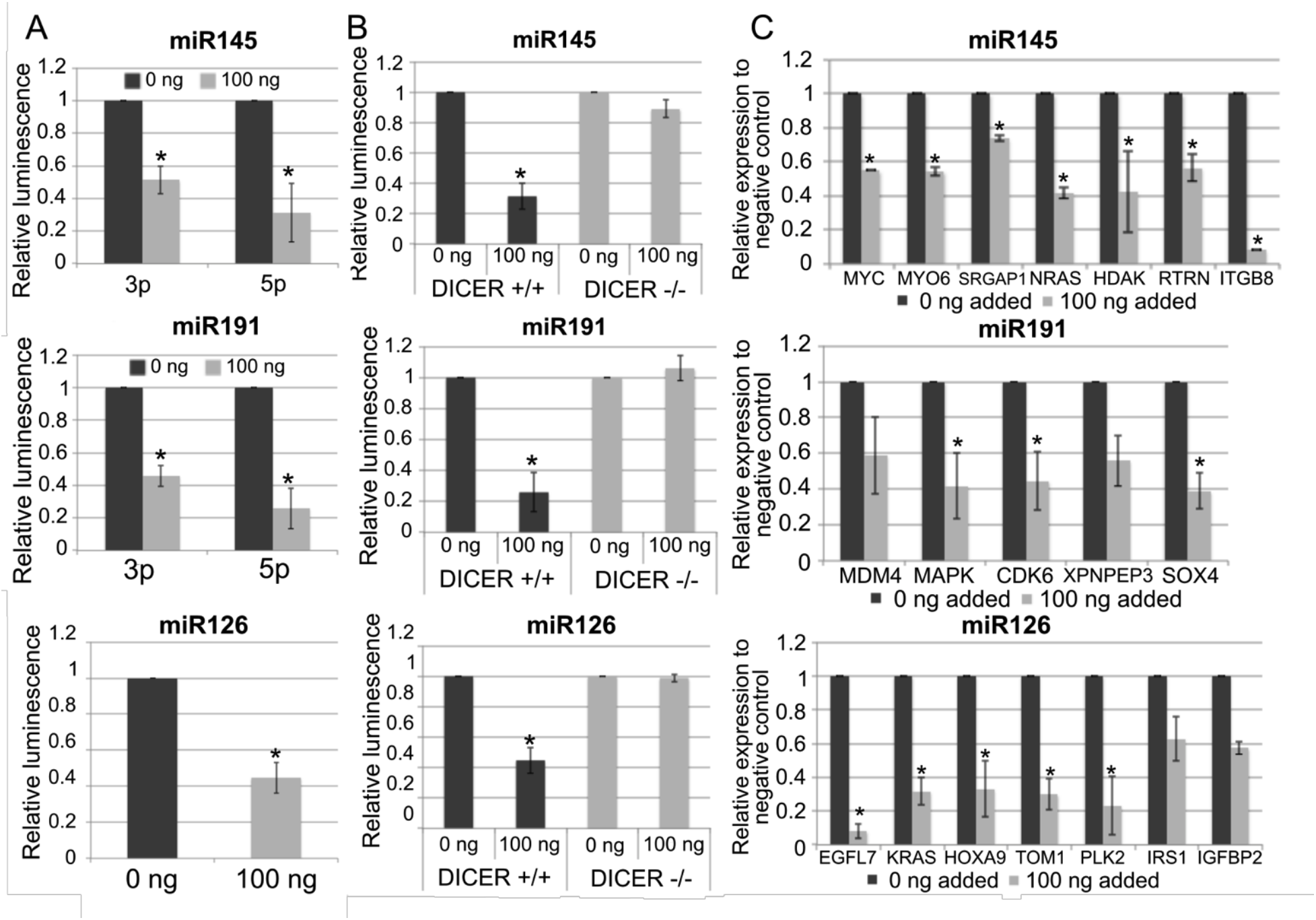
(A) Transfection of artificial microDNA, carrying indicated pre-miRNA sequences, to 293T cells decreases expression of a co-transfected *Renilla* luciferase reporter containing a sequence complementary to the miRNA sequence within its 3’ UTR. RL activity expressed relative to a cotransfected firefly luciferase and normalized again to the level in cells transfected with 0 ng of microDNA. Mean and S.E. of 3 experiments. * indicated p < 0.05 in a Student’s t test. (B) Repression of luciferase in dual-luciferase assay is observed in WT 293T cells but not in DICER1-/- 293T cells (C) Endogenous cellular targets of indicated miRNAs are repressed after transfection of the synthetic microDNA carrying the indicated pre-miRNA genes. mRNAs quantitated by Q-RT-PCR and expressed relative to the beta-actin gene and normalized to the level in cells transfected with 0 ng microDNA. Mean and S.E. of 3 experiments.

We next examined whether the repression of a luciferase reporter by a co-transfected microDNA is dependent on the endogenous RNA interference pathway. The introduction of each microDNA carrying microRNA sequences did not repress the luciferase reporter when transfected into *DICER1* KO 293T cells (35) (Figure 3B). This shows that the transcripts from microDNA need to be processed by Dicer through the same pathways as pre-miRNA produced from a chromosomal locus.

The microDNA carrying miRNA sequences also repress endogenous cellular genes that are targets of the encoded miRNA. The predicted targets were obtained from Targetscan. The microDNA carrying hsa-mir-145 sequence repressed mir-145 targets by up to 40%; hsa-mir-191 microDNA repressed mir-191 targets by up to 60%; hsa-mir-126 microDNA repressed downstream targets by up to 90% (Figure 3C). In each case most targets were repressed to similar levels. Further, the gene repression caused by a microDNA was specific to the targets of the miRNA encoded by the microDNA: for example, targets of miR-145 or miR-126 were not repressed by the introduction of microDNA containing miR-191 (Supplemental Figure 4C). Together these results show that the microDNA are potentially capable of contributing to the population of functional short RNA within a cell to influence the expression of endogenous genes.

### MicroDNA mimics containing exon sequences can repress host genes

MicroDNA are enriched from genes, and often contain exons (16). Short hairpin RNA (shRNA) sequences have long been known to be processed into siRNAs that repress target genes, suggesting that the gene from which a microDNA is derived could be repressed by the microDNA. Alternatively, if a microDNA is transcribed from both DNA strands the resulting double-stranded RNA could also be processed to a functional si-RNA (Figure 4A). Indeed, a similar phenomenon has been suggested in *Paramecium tetraurelia* where transposon-derived DNA sequences are ligated to form circles (of unknown size) that produce siRNAs that repress transposon expression (28).

**FIGURE 4.**
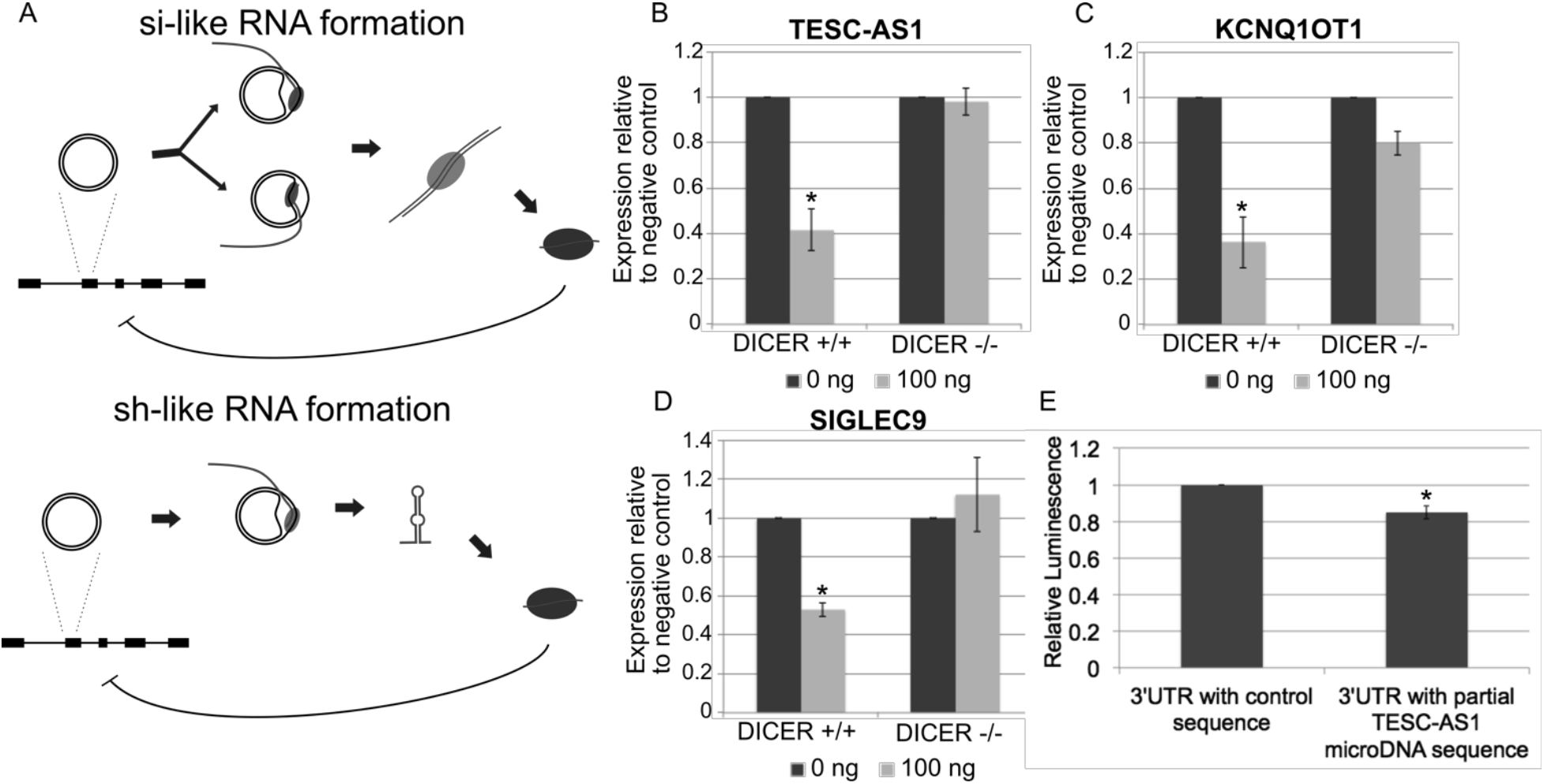
(A) Diagram of theoretical mechanism of the formation of si-like or sh-like RNA from microDNA containing an exonic sequence either from transcription of both strands of the microDNA or from folding of the transcript into a short-hairpin. (B) Transfection of a microDNA containing the sequence of exon 2 of the TESC-AS1 gene represses the expression of the TESC-AS1 gene in 293Tcells. The repression is observed in 293T cells but not in DICER-/- 293T cells. The same is seen with microDNA carrying a portion of (C) Exon 1 of KCNQ1OT1 and (D) the Exon 6 of SIGLEC9. The RNAs were quantitated by Q-RT-PCR and the values expressed relative to beta actin and normalized to the cells with 0 ng of transfected microDNA. Mean and S.E. of 3 experiments. *= P < 0.05 in a Student’s t-test. (E) A luciferase reporter assay where a sequence homologous to a known microDNA arising from the gene TESC-AS1 has been inserted into the 3’UTR of a *Renilla* luciferase gene. The luminescence of the *Renilla* luciferase gene is normalized against the luminescence from a synthetic firefly luciferase gene within the same plasmid.

To test whether microDNA could repress the parental gene, we created artificial microDNAs containing exonic sequences that we had identified in human cancer cells (Supplemental Table 1). These microDNA were transfected into 293T cells, where we get nearly 80% of the cells taking up the transfected DNA (data not shown). Three microDNA containing different exon sequences specifically repressed the expression of the host gene that contained the matching exon sequence. A microDNA carrying the full sequence of exon 2 of the *TESC-AS1* gene repressed the TESC-AS1 mRNA by ~60% (Figure 4B). A microDNA encoding an exon of the *KCNQ1OT1* gene repressed the KCNQ1OT1 mRNA by ~60% (Figure 4C). A microDNA encoding the full exon 6 of *SIGLEC9* gene repressed of the SIGLEC9 mRNA by ~50% (Figure 4D). Here again the repression of the endogenous gene by the microDNA is dependent on the RNA interference pathway, specifically *DICER1*, with the repression of TESC-AS1, KCNQ1OT1, or SIGLEC9 mRNAs significantly attenuated in *DICER1* KO 293T cells (Figure 4B-4D). The expression of genes not containing homology to the microDNA mimic were not affected by the introduction of microDNA: for example the microDNA from *KCNQ1OT1* did not repress *TESC-AS1* or *SIGLEC9* (Supplemental Figure 4D). This shows that the microDNA can also produce novel si-like RNA when the microDNA are derived from exonic sequences.

Thus regulatory short RNAs can be produced not only from microDNA which contain pre-microRNA genes but also from microDNA which overlap with exons. This greatly expands the proportion of eccDNA now expected to contribute to gene expression changes.

### Endogenous microDNA can repress a luciferase reporter

To test whether endogenous microDNA exist at high enough copy number to repress the expression of a gene, we transfected a luciferase reporter which contains within its 3’UTR a sequence homologous to a known endogenous microDNA from *TESC-AS1* (diagram of reporter shown in Supplemental Figure 5). The luciferase reporter carrying the sequence homologous to *TESC-AS1* was repressed to 80% relative to the control luciferase reporter with no *TESC-AS1* sequence even though no microDNA was co-transfected (Figure 4E). This suggests that regulatory short RNAs could arise from the endogenous *TESC-AS1* microDNA at a level sufficient to repress a transfected luciferase RNA containing the *TESC-AS1* sequence.

### Endogenous microDNA are associated with RNA polymerases

We next tested whether RNA polymerases bind to endogenous microDNA. Epitope-tagged (Halo Tag (HT)) POLR3F (subunit of RNA Polymerase III) or POLR2H (subunit of all three RNA Polymerases) were induced by doxycycline, in 293A cells (Figure 5). The epitope tagged subunits were then captured by covalent linkage to HaloLink resin and the pull-down confirmed by western blotting of the eluate for a non-covalently associated RNA polymerase subunit, POLR3A (Figure 5). The DNA which was associated with the HT alone or HT-POL subunits was isolated, digested by exonucleases, and then amplified by multiple displacement amplification with random hexamers (Figure 6A). The amplified DNA was quantitated and found to be significantly more in the POLR3F and POLR2H eluates than in the HT alone negative control precipitates: undetectable (below limit of detection) for the HT control, 728 ng for HT-POLR3F, and 408 ng for HT-POLR2H. Note that the HT alone is 33 kDa in size, and so a significant sized protein is being pulled down on the negative control HT beads. Further, PCR of the sheared RCA products ligated to sequencing adapters show that POLR3F and HT-POLR2H pull down 2e10 (or ~1000) fold more DNA than the HT control (Figure 6B). Because POLR2H and POLR3F both bind to endogenous microDNA, it suggests that microDNA could be transcribed by RNA Polymerase III, and possibly RNA Polymerase I and II.

**FIGURE 5.**
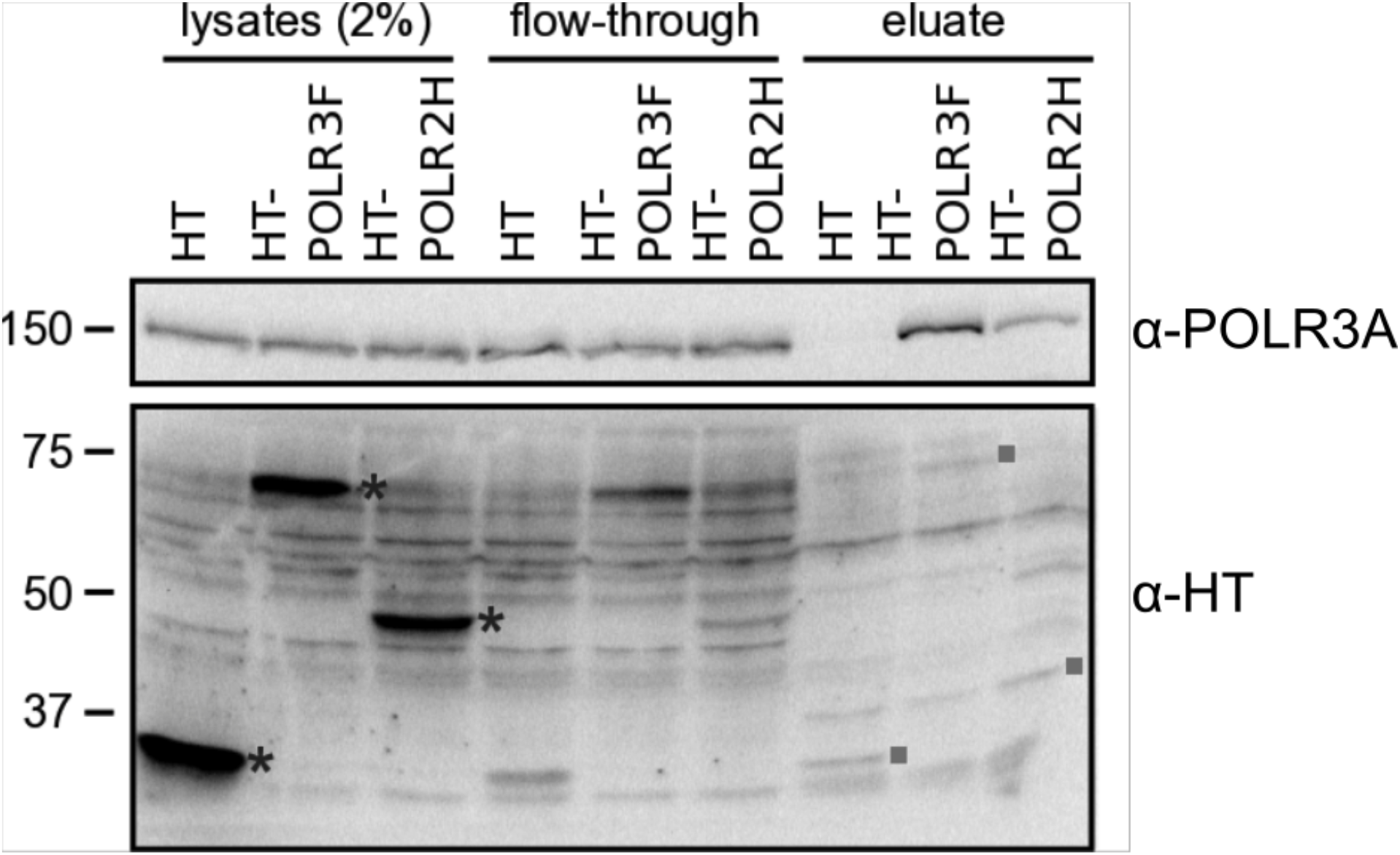
Efficiency of pull-down of RNA polymerase complex by HT. Detection of RNA polymerase III catalytic subunit (POLR3A) in the HT-POLR2H or HT-POLR3F pull-down. HaloTag fused RNA polymerase subunit POLR2H or POLR3F was expressed in 293A cells (asterisk). The HT-tagged polymerase subunits were covalently bound to the HaloLink resin and the non-covalently associated proteins were eluted by boiling in Laemmli Sample buffer. The HT-proteins are covalently bound to the beads and are mostly not eluted. Some covalent bonds break to release traces of the HT protein in the eluate (squares). However, non-covalently associated POLR3A of the RNA polymerase complex is specifically released in the eluates from the HT-POLR3F and HT-POLR2H pull-downs.

The rolling circle amplification products were subjected to Illumina sequencing to identify the microDNA and determine the number of unique sites in the genome represented in the microDNA library (complexity). The HT-POLR3F and HT-POLR2H associated libraries yielded 27-fold and 6-fold more complexity than the HT associated library (Table 1). Since the yield of DNA after rolling circle amplification was significantly more in the two POLR precipitates, we also compared the complexity by randomly sampling equal numbers of high-throughput reads from the three precipitates. Even after equalizing read numbers the HT-POLR3F and HT-POLR2H precipitates had microDNA at 3-8 fold higher complexity than that associated with just the HaloTag (Table 1 and Figure 6C), providing further evidence that microDNA bind to RNA polymerase subunits. Sequencing of the multiple displacement amplification products also confirmed that the microDNA associated with the RNA Polymerases have the same characteristics as found in our previous studies including length peaks at around 200-400 base pairs with a periodicity of nucleosome length DNA (Figure 6E), and GC content around 45-50% (Figure 6F). This shows that naturally occurring microDNA is associated with RNA polymerases, consistent with the hypothesis that they can be transcribed to produce functional regulatory small RNAs.

**FIGURE 6.**
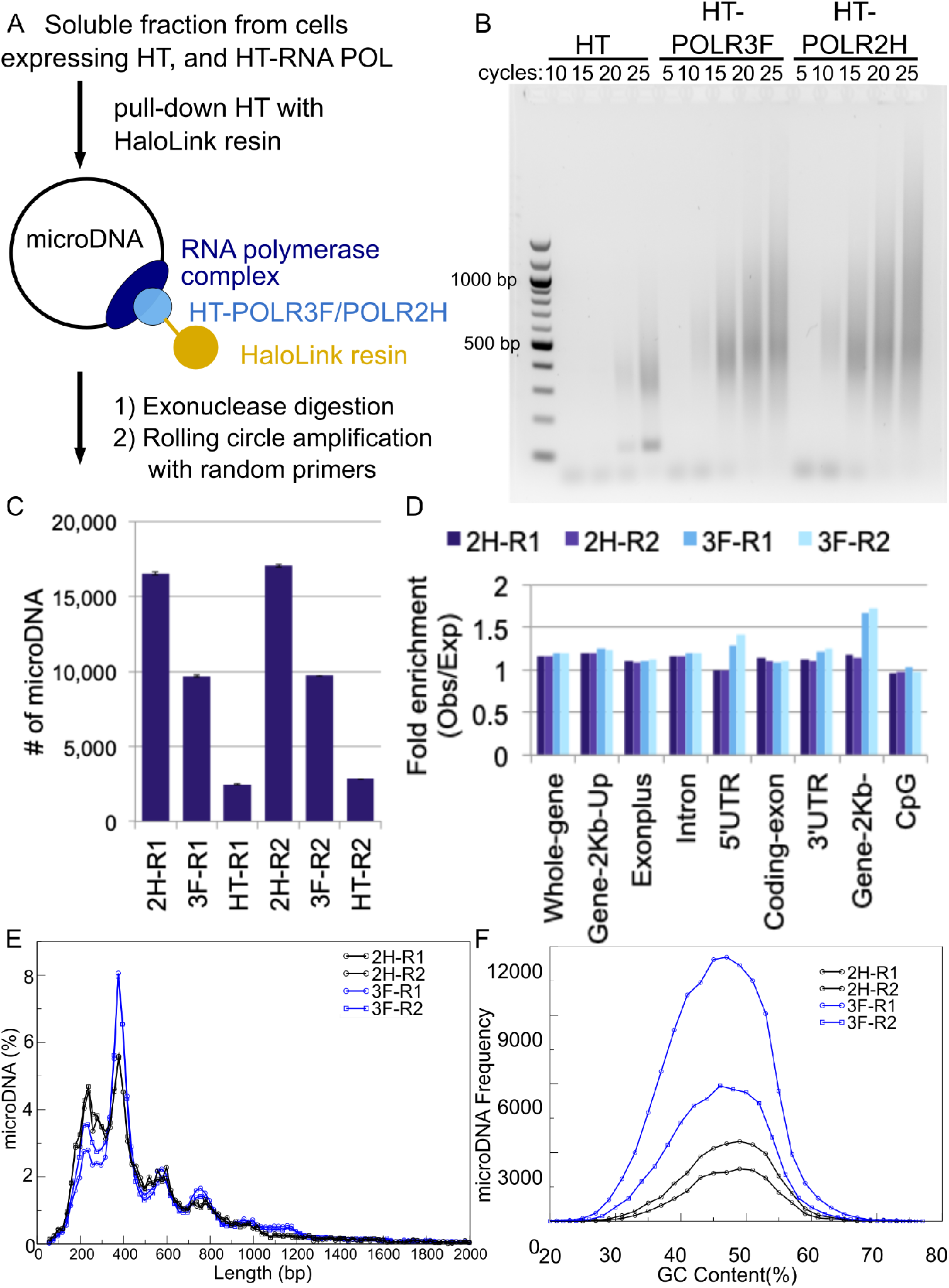
Subunits of PolI, PolII and PolIII bind to microDNA. (A) Diagram of pull-down of Halo-tagged RNA polymerase subunits and purification of associated microDNA for rolling circle amplification with random hexamers (RCA). (B) RCA products were sheared, ligated to library primers for high throughput sequencing, amplified by PCR for the indicated cycles and comparable aliquots run on a gel and visualized by ethidium bromide fluorescence. (C) Complexity of microDNA in the libraries prepared from the POLR3F and POLR2H pull-downs relative to the tag-only control as measured in Table 1. The error bars indicate the S.D. from 10 random subsamples from each library. (D-F) Characterization of microDNA molecules pulled down by POLR3F and POLR2H: (D) Enrichment relative to random expectation of the microDNA from areas of the genome with indicated genomic features, (E) length distribution, (F) GC content.

We have reported before that the microDNA in mammalian cells is 4-10X enriched relative to random expectation from 5’ UTRs, exons, genes, and CpG islands (16). In contrast, the RNA polymerase associated microDNA are more uniformly distributed throughout the genome (Figure 6D), suggesting that the RNA polymerase subunits have a lower affinity for microDNA from gene or CpG island derived areas.

## DISCUSSION

In summary, we have found that microDNA like molecules within cells are capable of being transcribed without a canonical promoter to form functional microRNA and novel sh-or si-RNA. The transcripts from microDNA mimics which carry microRNA sequences are processed into mature microRNA which can repress expression of down-stream targets and a luciferase reporter. MicroDNA arise from about 4% of the genome in the chicken DT40 cell line and 0.4-1% of human HeLa, C4-2, and LNCap cell lines (Supplemental Figure 6), and are enriched from genic regions (13,14,16). An intriguing finding of this study is that microDNA mimics which carry exonic sequences form novel sh-or si-RNA that repress the gene from which it originated. Further, a luciferase reporter carrying a sequence homologous to an endogenous microDNA sequence is repressed when transfected into human cancer cells without any co-transfected synthetic microDNA, suggesting that sh-or si-RNA from endogenous microDNA is present at a level sufficient to be functional. Consistent with this, subunits within RNA polymerase complexes (POL2H and POL3F) are bound to naturally occurring microDNA, adding to the possibility that endogenous microDNA may be transcribed. Collectively, these results support the hypothesis that microDNA could be functional in cells, actively repressing genes through the RNA interference pathway by producing microRNA and novel si-RNA.

EccDNA, including microDNA, are found to be significantly increased in cancer cells (13,14,16,21). We know that long eccDNA like Double Minutes can amplify genes, including oncogenes, in cancer and change gene expression patterns that contribute to oncogenesis. Our current results suggest that the microDNA could similarly regulate gene expression patterns through the formation of novel regulatory RNAs arising from microDNA.

Interestingly, it has recently been discovered that artificial eccDNA molecules introduced into cancer cell lines have high stability but are quickly transcriptionally silenced by epigenetic mechanisms (36). Although 359 bp long microDNA like plasmids have been shown to be assembled *in vitro* into mono-or di-nucleosomes (37), it is unclear whether the microDNA in cells are chromatinized, and if they are, whether they can be epigenetically repressed like the long eccDNA. Therefore, it will be interesting to test in the future whether the microDNA can or cannot be epigenetically regulated.

More research is necessary to determine the exact mechanism by which microDNA are transcribed, but it is clear that the circularization is important, because the equivalent linear DNA was not transcribed *in vitro* (Figure 1C and Supplemental Figure 2) or *in vivo* (Supplemental Figure 4A). We believe that the transcription of microDNA without the requirement of a canonical promoter sequence is possible because of structural features unique to the microDNA. At most 1 or 2 nucleosomes could be assembled on circles of 359 bp *in vitro* (37,38). Thus abnormal chromatinization may leave the intervening naked DNA more accessible to transcription factors including RNA polymerases (39) allowing promoters or cryptic promoters to be more readily bound. The small size of the microDNA molecule may also contribute to its spontaneous transcription through the formation of flipped bases (40) and bubbles of ssDNA (41), the latter known to initiate transcription (42). Further, the bent shape of the small circular DNA itself may signal for the binding of TATA-binding proteins (31–34), though the bent shape of DNA can also limit its transcription when associated with a *bona fide* promoter (43). Lastly, it has also been shown that nicks can initiate transcription which suggests that if microDNA molecules contain nicks then that could also contribute to their transcription (44).

We have previously reported that microDNA arise from epigenetically active gene-rich areas within the genome (16). Therefore, transcription can lead to microDNA formation, which could lead to repression of the parent gene by generating regulatory short RNAs. This could be a negative feedback mechanism that represses GC-rich genes when they are transcribed at a high level and produce microDNA, most likely from some DNA repair process. It is intriguing that a similar pathway has been proposed for the silencing of transposon genes in the germline nucleus of *Paramecium tetraurelia* where transposon-derived DNA sequences are ligated to form circles (of unknown size) that produce siRNAs that associate with PIWI proteins to repress transposon expression (28).

One criticism of our study is that the amount of synthetic microDNA we transfected into cells to see increase in microDNA-encoded regulatory RNAs is significantly higher than endogenous levels of microDNA. We believe that this is because of inefficiencies introduced by several factors: the presence of endogenous microRNA that sets a high basal level of the RNA for comparison, the low number of DNA molecules taken up per cell and the possible saturation of endogenous RNA polymerases. The microDNA sequences selected for this study were chosen because of the existing knowledge of their encoded microRNAs and their target genes. As a result, the fold-increase of microRNAs arising from transfected microDNA is diminished by the high basal level of endogenous microRNA in the untransfected cell. Indeed, when we look for microDNA-junction-specific RNAs, the fold-change is very high because there is very little endogenous RNA to set a high basal level (Supplemental Figure 4B). Another important factor is that we do not know how much of the added microDNA is being taken up by the cells. It is very likely that these fractions/numbers are low leading to an underrepresentation of the effect of the synthetic exogenous microDNA. In addition, the microDNA molecules have no inherent structural feature that would cause them to escape from the endo-phagosome and be transported into the nucleus therefore limiting the amounts of the transfected microDNA that can be transcribed. Finally, it is unknown if there is sufficient free RNA polymerase to associate with the newly arriving nuclear microDNA, further decreasing the proportion of exogenous microDNA which is actively transcribed. Overall, we believe the effect of the microDNA transfected into the cells is underrepresented by our experiments because of these limiting factors.

In summary, inspired by the prevalence of small eccDNA (microDNA) in normal and cancer cells, we examined whether molecules with similar characteristics could express functional gene products. The results suggest that microDNA could express regulatory short RNAs, raising the possibility that microDNA could cause changes in cell phenotype by regulating gene expression.

## ACKNOWLEDGMENTS

We thank all members of the Dutta lab for useful discussions. This work was supported by NIH R01s CA166054 and CA60499 to AD.

