## Supplemental Materials for "Extrachromosomal circular DNA, microDNA, without canonical promoters produce short regulatory RNAs that suppress gene expression"

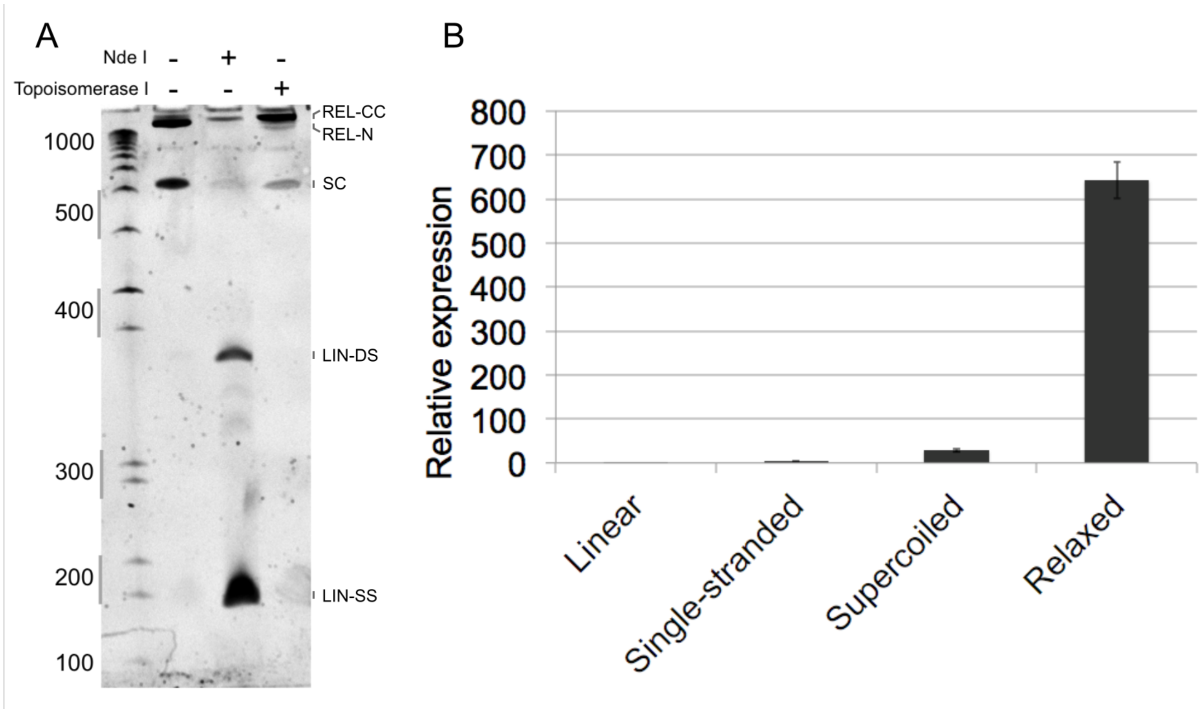


**Supplementary Figure 1** Circular microDNA mimic molecules of all topologies are transcribed by RNA polymerases in HeLa nuclear extract *in vitro* transcription assay. (A) Different topologies of microDNA (with KCNQ1OT1 sequence as example) analyzed by incubation with NdeI digestion (cuts the double-stranded microDNA twice) or topoisomerase (relaxes supercoiled microDNA). REL-CC: Relaxed covalently closed circular DNA; REL-N: Relaxed nicked circular DNA; SC: Supercoiled circular DNA; LIN-DS: Linear double stranded DNA; LIN-SS: Linear single stranded DNA. Representative replicate of duplicates. (B) QPCR of RNA products obtained by IVT of microDNA with indicated topologies or strandedness (with hsa-mir-191 sequence) Mean and S.D. of 3 replicates.

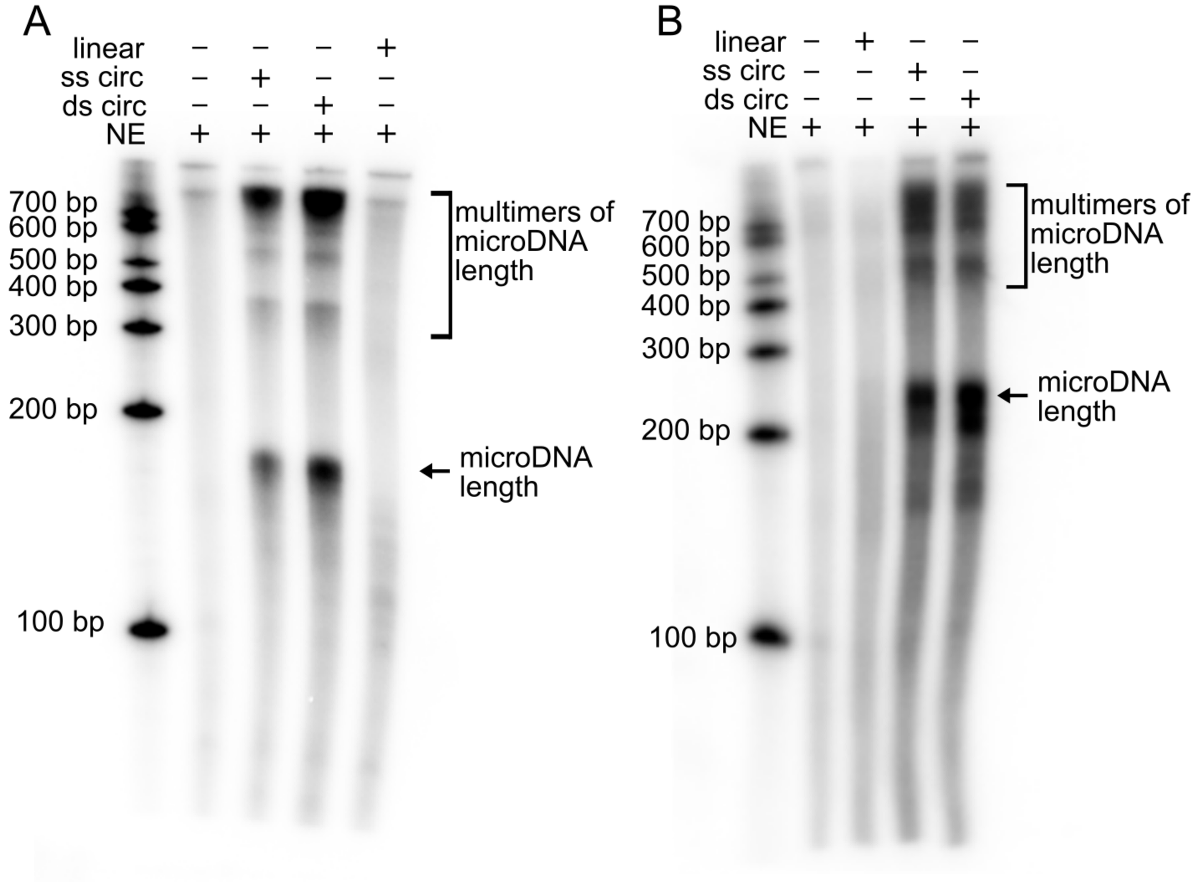


**Supplementary Figure 2** The in vitro transcription of microDNA by RNA polymerases within HeLa Nuclear Extract is verified with two other microDNA sequences: (A) microDNA containing hsa-let-7a (B) microDNA containing hsa-mir-145. Representative replicate of duplicates.

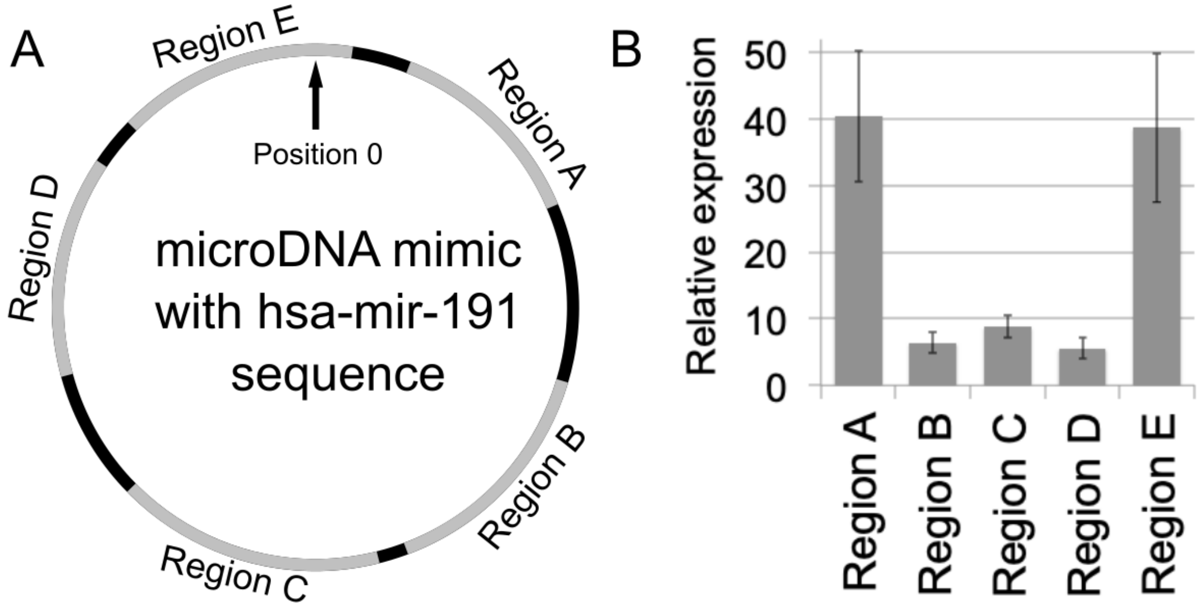


C DNA sequence of each region:

A:TTTAGCGAGATGCGGGCTGAGGCAGCCCTGGGGGAGTCACTACCATTGC

B:CTCCCCAGGGTTCCAGAGATGGCCACCAGCCTCGGCTCTCCCACGTCTACAC

C:CCTGAGGGCGCAACTGAGAGGAACTGAGACCCAAGCAGCTCAGTAAAGTATGTC

D:AGGAGAGCAGGGGACGAAATCCAAGCGCAGCTGGAATGCTCTGGAGACAACAG

E:TTCCGTTGCCCGCTGTCCAGCCGTTGGCGGGGGGCGGGGGAGCTCCTGCCCCC

**Supplementary Figure 3** Distribution of transcripts from microDNA is not random (A) Diagram of regions of the microDNA whose transcripts are quantitated by Q-RT-PCR. (B) Quantification of RNA arising from each region of the microDNA after normalizing to efficiency of the primer pairs when they are used to amplify the circular DNA. Mean and S.E. of 3 replicates. (C) Sequence of microDNA within each specified region

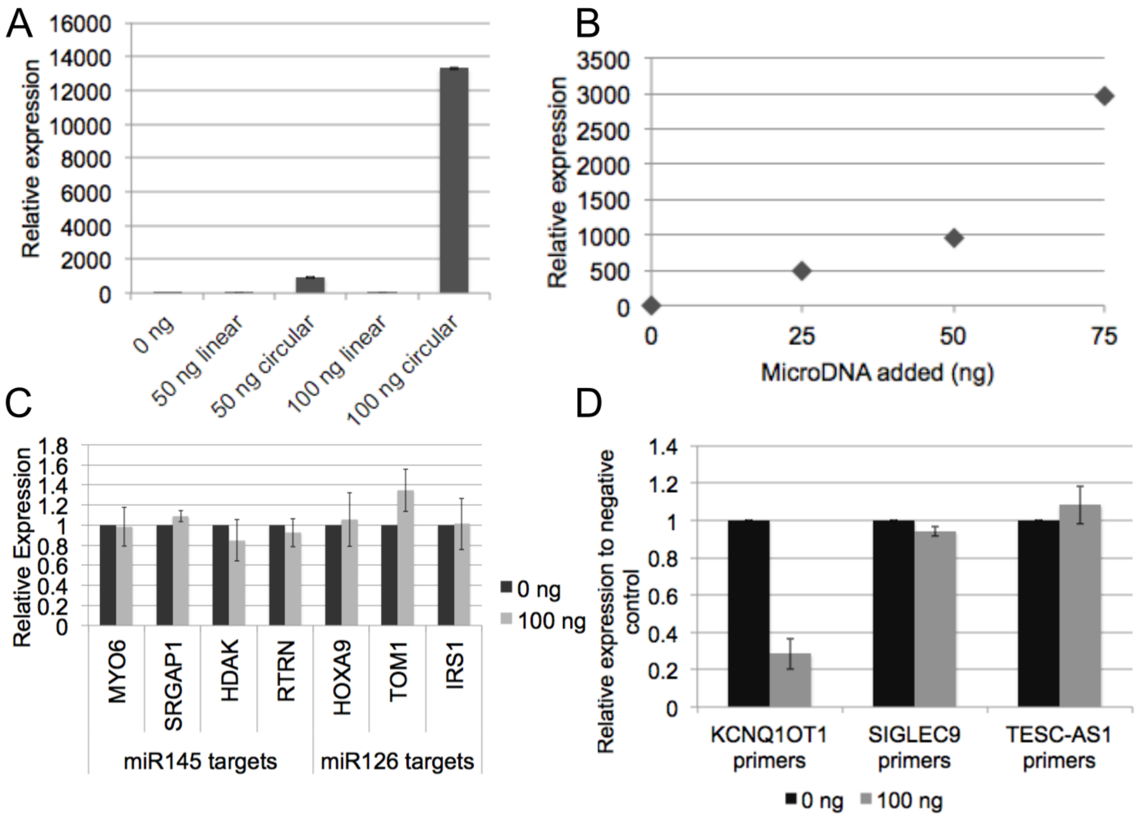


**Supplementary Figure 4** RNA transcribed from circular microDNA specifically and the repression of gene expression by RNA transcripts arising from microDNA is sequence-specific. (A) Circular, but not linear, molecules of the hsa-mir-126 microDNA increases the encoded RNA when transfected into 293T cancer cells. Mean and S.D. of 3 replicates. (B) QPCR of RNA arising only from microDNA quantified using primers which amplify the junction sequence of the microDNA (containing hsa-mir-126) Mean and S.D. of 3 replicates. (C) When the microDNA containing the miR-191 sequence was transfected into human cancer cells, genes which are not targeted by miR-191 were not repressed. Mean and S.D. of 3 replicates. (D) When the microDNA containing the KCNQ1OT1 microDNA sequence was transfected into human cancer cells, genes other than KCNQ1OT1 were not repressed. Mean and S.D. of 3 replicates.

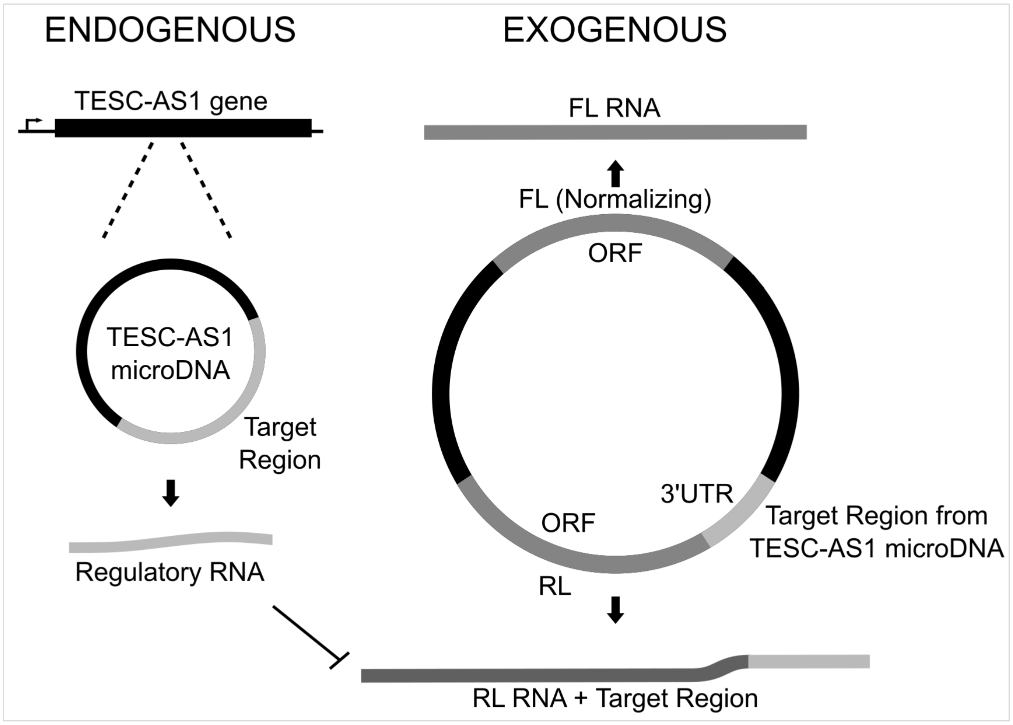


**Supplementary Figure 5** Diagram of the luciferase reporter assay used to test whether endogenous microDNA have the ability to repress gene expression. The luciferase reporter was transfected into a human cancer cell line and the repression of the *Renilla* luciferase gene with the TESC-AS1 homologous sequence was normalized to the control luciferase gene (the synthetic firefly luciferase gene).

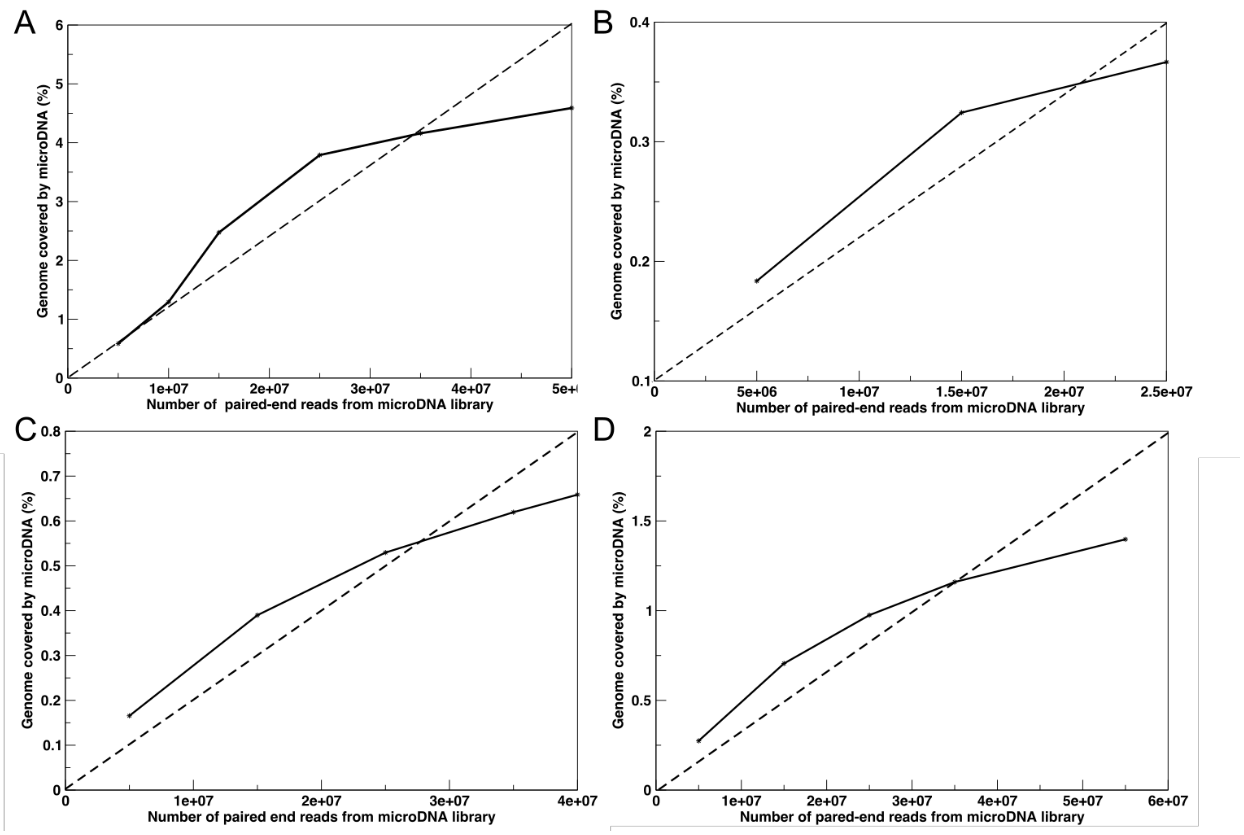


**Supplementary Figure 6** Percentage of the genome that give rise to microDNA in (A) DT40, (B) HeLa, (C) C4-2, and (D) LNCaP cells. As we prepare many libraries and sequence them, the % of the genome covered by the microDNA sequences does not increase linearly but appears to saturate at 4-5 % of the genome for chicken DT40 cells, and 0.4-1.5% of the genome for human HeLa, C4-2, and LNCaP cells.

| Name | | | Primer Sequence | MicroDNA sequence | Legend |
|---|---|---|---|---|---|
| miR145 | AB | F | CCCCCCAGAGCAATAAGCC | CCCCCCAGAGCAATAAGCCACATCCGGCGACGTGTGGCACCCCACCCTGG<br>CTGCTACAGATGGGGCTGGATGCAGAAGAGAACTCCAGCTGGTCCTTAGG<br>GACACGGCGGCCTTGGCGCTGAAGGCCACTCGCTCC**CACCTTGTCCTCA**<br>**CGGTCCAGTTTTCCCAGGAATCCCTTAGATGCTAAGATGGGGATTCCTGG**<br>**AAATACTGTTCTTGAGGTCATGGTT**TCACAGCTGGATTTGCCTCCTTC | Bold: pre-hsa-mir-145 sequence<br>Underlined: 5p and 3p miR145 sequence |
| | | R | GAAGGAGGCAAATCCAGCTGT | | |
| | CD | F | GGCCACTCGCTCCCACCTTG | | |
| | | R | TTCAGCGCCAAGGCCGC | | |
| miR191 | AB | F | CCCCAGGAAGTAAGAGGGCTATCTTTAGCG | GCAGGAGCTCCCCCGCCCCCCGCCAA**CGGCTGGACAGCGGGCAACGGA**<br>**ATCCCAAAAGCAGCTGTTGTCTCCAGAGCATTCCAGCTGCGCTTGGATTT**<br>**CGTCCCCTGCTCTCCTGCCT**GAGCAGCGCCCTGGCCCAGATGGGGTGCC<br>CCTGACCCCCAGACATACTTTACTGAGCTGCTTGGGTCTCAGTTCCTCTCA<br>GTTGCGCCCTCAGGCTGGAGGTGATGGGTGTAGACGTGGGAGAGCCGAG<br>GCTGGTGGCCATCTCTGGAACCCTGGGGAGGATTGGCGAGGGAGGGTGG<br>ACCCAGGACCTCTGGGTAGGGCTGCAATGGTAGTGACTCCCCCAGGGCTG<br>CCTCAGCCCGCATCTCGCTAAAGATAGCCCTCTTACTTCCTGGGG | Bold: pre-hsa-mir-191 sequence<br>Underlined: 5p and 3p miR191 sequence |
| | | R | GCAGGAGCTCCCCCGCCC | | |
| | CD | F | AGAGGAACTGAGACCCAAGCAGC | | |
| | | R | CAGTTGCGCCCTCAGGCT | | |
| miR126 | AB | F | GGAGGATAGGTGGGTTCCCGA | GGAGGATAGGTGGGTTCCCGAGAACTGGGGGCAGGTTGCCCGGAGCCTC<br>ATATCAGCCAAGAAGGCAGAAGTGCCCCGTCCCGGGGTCCTGTCTGCATC<br>CAGCGCAGCATTCTGGAAGACGCCACGCCTC**CGCTGGCGACGGGACATT**<br>**ATTACTTTTGGTACGCGCTGTGACACTTCAAACTCGTACCGTGAGTAATAA**<br>**TGCGCCGTCCACGGCA**CCG | Bold: pre-hsa-mir-126 sequence<br>Underlined: 5p and 3p miR126 sequence |
| | | R | CGGTGCCGTGGACGGCGCATT | | |
| | CD | F | TTCTGGAAGACGCCACGCCTC | | |
| | | R | TGCTGCGCTGGATGCAGACAG | | |
| Let-7 | AB | F | TCCTCAGCCCTCTTTCCTCC | TCCTCAGCCCTCTTTCCTCCCGCGTCCCCAGGAGGTGCCTCTGGAAGCCA<br>CGGAGTCCCATCGGCACCAAGACCGACTGCCCTTTG**GGGTGAGGTAGTA**<br>**GGTTGTATAGTTTGGGGCTCTGCCCTGCTATGGGATAACTATACAATCTACT**<br>**GTCTTTCCT**GAAGTGGCTGTAATATCTGCGGTG | Bold: pre-hsa-let-7a sequence<br>Underlined: 5p and 3p let7a sequence |
| | | R | CACCGCAGATATTACAGCCACTTC | | |
| | CD | F | GGTAGTAGGTTGTATAGTTTGGG | | |
| | | R | CCAAAGGGCAGTCGGTCTT | | |
| TESC-AS1 | AB | F | TCTCCATCCAGTTGGGGTGACT | TCTCCATCCAGTTGGGGTGACTGGGGCCGATGCTGTTGCCTGTGTAATGTG<br>ATTCCTCCTCCTTAAAATAAG**GATAAGTTAATGAGATGTCCACCAGAGGGC**<br>**CTGGCGTGGACACAGCACATGGCCACAGAGGCTGGTGGGCGCTATGGAA**<br>**TCTTGTCCCCTGGAGAGGCACACAGCCAGGGCAGAACATCAAGGTCAAG**<br>**GCTCTCCTGAAGGCTCTGCAGTGCTTAGTGACACCACCATCAATGACCGT**<br>**CAGGTATCAG**GTCTGTTCTGAACGTTAGTCT | Bold: Portion of TESC-AS1 exon 2 |
| | | R | AGACTAACGTTCAGAACAGACCTGA | | |
| | CD | F | CCCACCAGCCTCTGTGGC | | |
| | | R | CGCTATGGAATCTTGTCCCCTG | | |
| SIGLEC9 | AB | F | CAGGCTACATGCTGGCTGTG | CAGGCTACATGCTGGCTGTGGAGAGTCCACATCACTCAC**CTGAGAGGCTG**<br>**AACCCCTGACAGCGTTTGCATCCTCTATGCCCGTATCTCCCACGCCCGCT**<br>**GCTGGCCTTGCCGATTTCTTCCTGCAGGACCTCACT**CTGAGTGAAGAGAC<br>CAGAGAGCCTTTCAGTGTGGTCAGATTGG | Bold: Portion of SIGLEC9 exon 6 |
| | | R | CCAATCTGACCACACTGAAAGG | | |
| | CD | F | CACGCCCGCTGCTGGCCT | | |
| | | R | GGAGATACGGGCATAGAGGATG | | |
| KCNQ1OT1 | AB | F | GAAAGGACACCATGTTGAACACATC | **ATCCCTTGTCAGATGAATATTTTCTCCCATTCTACAGGATATCTCTTAATAGT**<br>**TTATTATTTCCTTTTATGTGTAGGTTTTTAGTTTGATACAGTCCCATTTGTCTA**<br>**TTTTTGTTTTTGTTGCCTGTGCTTATAAAGTCTTACCCATAAAATCTTCGCTA**<br>**GACCCATATCCTGAAGGGTTTCCCTTCTGTTTTCTTCTAGTACTGTTTGCTT**<br>**CAGGTCTCATGTTTAAGTCTCTAATCAATTTTGAGTTGATTTTTTTATATGCT**<br>**GAGAGATAGTATAGTTTCATTCTTCTGCATATGATATCCTGTTTTCCCAACAT**<br>**GATGTGTTCAACATGGTGTCCTTTC** | Bold: Portion of KCNQ1OT1 exon 1 |
| | | R | ATCCCTTGTCAGATGAATATTTTCTCC | | |
| | CD | F | AGGATATGGGTCTAGCGAAGATTTTATG | | |
| | | R | GAAGGGTTTCCCTTCTGTTTTCTTCTA | | |

**Supplementary Table 1** Artificial microDNA were created using sequences of endogenous microDNA found in previous studies (Shibata 2012, Dillon 2015) which overlapped with microRNA.

The primer sequences amplified the staggered duplexes AB or CD for each microDNA. When these duplexes were annealed and subjected to LAMA we obtained the circular microDNA derived entirely from genomic sequence which matched the microDNA sequence acquired in previous studies. The sequences are listed under the 'MicroDNA sequence' column.
